# Synergy mediates Long-Range Correlations in the Visual Cortex Near Criticality

**DOI:** 10.1101/2025.11.07.687179

**Authors:** Hardik Rajpal, Cedric Stefens, Meghdad Saeedian, Joe S. Canzano, Michael G. Kareithi, Mauricio Barahona, Spencer LaVere Smith, Simon R Schultz, Henrik Jeldtoft Jensen

## Abstract

Long-range correlations are a key signature of systems operating near criticality, indicating spatially-extended interactions across large distances. These extended dependencies underlie other emergent properties of critical dynamics, such as high susceptibility and multi-scale coordination. In the brain, along with other signatures of criticality, long-range correlations have been observed across various spatial scales, suggesting that the brain may operate near a critical point to optimise information processing and adaptability. However, the mechanisms underlying these long-range correlations remain poorly understood. Here, we investigate the role of synergistic interactions in mediating long-range correlations in the visual cortex of awake mice. We leverage recent advances in mesoscale two-photon calcium imaging to analyse the activity of thousands of neurons across a wide field of view, allowing us to confirm the presence of long-range correlations at the level of neuronal populations. By applying the Partial Information Decomposition (PID) framework, we decompose the correlations into synergistic and redundant information interactions. Our results reveal that the increase in long-range correlations during visual stimulation is accompanied by a significant increase in synergistic rather than redundant interactions among neurons. Furthermore, we analyse a combined network formed by the union of synergistic and redundant interaction networks, and find that both types of interactions complement each other to facilitate efficient information processing across long distances. This complementarity is further enhanced during the visual stimulation. These findings provide new insights into the computational mechanisms that give rise to long-range correlations in neural systems and highlight the importance of considering different types of information interactions in understanding correlations in the brain.

## 1. Introduction

The theory of self-organised criticality (SOC) provides a framework for understanding how complexity arises in natural systems and a balance between order and disorder is maintained^1,2^. SOC suggests that complex systems naturally evolve towards a critical state, and, in such a state, small perturbations can lead to large-scale events, often characterised by power-law distributions^3^. This critical state exhibits enhanced information processing^4–6^ and computational capabilities^7^, as it allows for a balance between stability and adaptability^4–6^. In addition, systems at criticality exhibit other desirable properties, such as maximal dynamic range^8^ and long-range correlations across regions that can facilitate information integration ^9^. These properties have inspired the *brain criticality hypothesis*^10–12^, which posits that the brain operates near a critical point to optimally process information, respond to a wide range of stimuli, and orchestrate a balance between functional segregation and integration^13–15^.

Empirical evidence supporting the brain criticality hypothesis has been observed across various spatial and temporal scales, from the microscopic level of individual neurons to the macroscopic level of large-scale brain networks. At the microscopic level, studies have reported that neuronal avalanches, which are cascades of neuronal firings separated by silence, follow power-law distributions, in accordance with the critical brain hypothesis^16,17^. At the macroscopic level, functional magnetic resonance imaging (fMRI) studies have shown that large-scale brain networks exhibit scale-free dynamics and long-range correlations, which are also suggestive of a critical state^18,19^.

More recently, studies have explored the role of criticality in cognitive processes, such as sensorimotor processing, perception, attention, and memory^20–23^. Some studies have found that different signatures of brain criticality are sensitive to various states of consciousness, such as sleep, anaesthesia, and disorders of consciousness^24–26^. In computer science, it has been shown that artificial neural networks (ANNs) can benefit from operating near criticality, as it enhances generalisation performance and enables learning of optimal representations^27–30^. See recent reviews for a comprehensive overview of the state of research on brain criticality and its implications for brain dynamics in health and disease^6,10–15,31^.

Despite the growing body of evidence supporting the brain criticality hypothesis, several challenges and open questions remain. Among the various signatures of criticality, the correlation length, which quantifies the spatial extent of neural correlations, is expected to diverge at criticality^32^. Functionally, the ensuing long-range correlations are crucial for facilitating coordination among various brain regions and information integration. While some work has been done to explore the presence of long-range correlations in macroscopic scales in fMRI studies^18^, there is a need for experimental and computational studies to understand how long-range correlations manifest at the level of neuronal populations and how they relate to different cognitive states. So far, such studies have been limited to either in-vitro neuronal cultures or smaller populations of in-vivo neurons due to a lack of large-scale recordings.

Beyond identifying long-range correlations, it is also important to explore biological or computational mechanisms that give rise to spatially extended interactions. To address these questions, however, one needs to further decompose the nature of correlations in neural systems. Information theory provides a powerful framework to decompose the interdependencies between components of a system into different types, such as redundant, unique, and synergistic^33^. Previous work explored how different types of information interactions emerge from stimulus-evoked *vs*. stimulus-independent correlations^34,35^. Recent developments in information decomposition, such as the Partial Information Decomposition (PID)^33^ and the Integrated Information Decomposition (ϕ ID)^36^, have provided new tools to dissect the nature of correlations in neural systems. These approaches allow us to quantify the amount of redundant or synergistic information between different components of a system, as well as their unique contributions to a target variable or to the future state of the system. Synergistic interactions are particularly interesting, as they represent the information that is only available when considering multiple components together and cannot be obtained from any single component alone. For example, a post-synaptic neuron firing as an *XOR* gate of the inputs of the pre-synaptic neurons is a purely synergistic interaction, as the state of each pre-synaptic neuron alone does not provide any information about the state of the post-synaptic neuron^36^. Beyond neuroscience, synergy has been used to understand the different musical styles of Bach and Corelli^37^; to differentiate the technological complexity of economies^38^; and to study collective behaviour of cells in organoids^39^.

In this study, we leverage recent advances in mesoscale two-photon calcium imaging^40^ that make it possible to record the activity of thousands of neurons across a wide field of view (up to several millimetres) of the visual cortex of awake mice. This allows us to explore the presence of long-range correlations and their relationship to criticality in neuronal populations. We then apply the PID framework to decompose the correlations into different types of information interactions and investigate how these interactions vary across different spatial scales. By combining large-scale in-vivo neuronal recordings with advanced information-theoretic analysis, we aim to provide computational insights into the mechanisms that give rise to long-range correlations. We also explore how these correlations and information interactions vary between spontaneous and visually stimulated states.

We find that the visual cortex exhibits long-range correlations that extend across several millimetres, and these correlations are enhanced under visual stimulation. By applying PID, we find that synergistic information interactions play a crucial role in mediating long-range correlations. Indeed, redundant interactions are dominant and have a longer correlation length; synergistic interactions exhibit a more pronounced increase at large distances during visual stimulation. We further analyse a combined network constructed from the synergistic and redundant interaction layers, and find that both synergistic and redundant interactions complement each other to facilitate information processing across long distances. Our findings provide further support towards the brain criticality hypothesis by characterising long-range correlations in the visual cortex. Further, long-range correlations are preferentially modulated by synergistic interactions among the neurons under visual stimulation. These results provide novel insights into the role of synergistic interactions in the brain to coordinate activity across brain regions and their relationship to the possible critical brain dynamics. Furthermore, it highlights the importance of considering different types of information interactions in understanding neural systems.

## 2 Materials and Methods

The study is based on calcium imaging data recorded from the posterior cortex of awake mice using a two-photon mesoscope, including several visual areas. The datasets were preprocessed to extract the neural activity traces, which were then used for information-theoretic analyses. In this section, we describe the data acquisition, preprocessing steps, and the information-theoretic measures employed in our analysis.

### 2.1 Data Acquisition

All experimental procedures were approved by the Institutional Animal Care and Use Committee (IACUC) at the University of California, Santa Barbara and were conducted in accordance with the guidelines of the US Department of Health and Human Services. The animals used in the study were adult, triple transgenic mice of the genotype TITL-GCaMP6s (Ai94,^41^) X Emx1-Cre (Jackson Labs #005628) X ROSA:LNL:tTA (Jackson Labs #011008) to express the calcium indicator GCaMP6s in cortical excitatory neurons. The mice were housed in a 12-hour reverse light/dark cycle and had *ad libitum* access to food and water. A total of 10 recordings (5 spontaneous and 5 stimulated) were obtained from 8 mice, with some mice contributing multiple recordings on different days. At least one week before experiments, mice were surgically implanted with a 5mm optical glass coverslip over the right posterior cortex and a stainless steel headplate, both adhered with cyanoacrylate glue (Oasis Medical) and dental cement (Parkell Metabond), as previously described^42^. After recovery, intrinsic signals optical imaging (**ISOI**)^43^ was performed to measure cortical retinotopic maps used to delineate visual area boundaries, also as described previously^4445^. These were used to validate craniotomy targeting and to register the two-photon fields of view to visual areas via vascular landmarks.

The calcium imaging datasets were recorded at the University of California, Santa Barbara, using a custom-built two-photon mesoscope (Diesel2p)^40^. The mesoscope allows for imaging large fields of view (herein, 3 mm *×* 3-4 mm) at resonant speeds and cellular resolution. The imaging was performed at a median frame rate of 7.5 Hz and 1.46 microns per pixel. Fields of view were positioned to cover primary visual cortex (V1) and several higher visual areas (HVAs) at once, at depths of 150-200 *μ*m to capture L2/3 cortical neurons. Imaging sessions lasted approximately 45 minutes, during which the mice were either in a spontaneous state (no visual stimulus) or were presented with drifting grating patches as visual stimuli. The visual stimuli were presented on a 90 cm curved monitor placed 14.5 cm from the mouse’s left eye and tilted 10 degrees downward to cover approximately 150 degrees of visual angle in azimuth and 70 degrees in elevation. Drifting grating patches were presented sequentially as 20-degree-wide squares tiling the visual field area above. Each had a spatial frequency of either 0.04 or 0.1 cpd and a temporal frequency of 1 or 4 Hz, and were presented in 8 different orientations (0, 45, 90, 135, 180, 225, 270, and 315 degrees). The parameters of the patches in each sequence were randomly shuffled across successive presentations to enforce an even stimulus distribution. Finally, a projective correction was applied to screen stimuli to account for the near placement of a flat screen and the spherical eye of the mouse^46^; with the correction applied, images strictly maintain the intended geometry and location across the visual field relative to the eye centre. Visible emission from the screen was blocked from the imaging objective using a custom opaque plastic cone placed between the headplate and objective. Spontaneous recordings were obtained in the absence of any visual stimuli, with the mice either in total darkness or with the same screen set to a static middle grey image.

### 2.2 Preprocessing

The raw calcium imaging data were first preprocessed using the Suite2p pipeline^47^, which includes motion correction and alignment using the registration module. The motion correction step in Suite2p involved aligning the frames of the imaging data to correct for any movement artefacts. The cell detection and extraction of the fluorescence traces were performed using the EXTRACT algorithm^48,49^, which identifies regions of interest (ROIs) corresponding to individual neurons and extracts their fluorescence signals, while correcting for any neuropil contamination. The extracted fluorescence traces were then deconvolved to estimate the underlying spiking rate using the deep-learning-based CASCADE algorithm^50^. Finally, to obtain the discretized binary states of each neuron, the deconvolved traces were fitted using a Hidden Markov Model (HMM) to identify the most probable state (Active: 1 or Quiet: 0) of each neuron at each time point (see Figure 1). This approach allows us to avoid setting an arbitrary threshold for each neuron and provides a probabilistic framework for identifying discrete states from the deconvolved calcium traces^51,52^. The binarised states were then used for the correlation and information-theoretic analyses.

**Figure 1.**
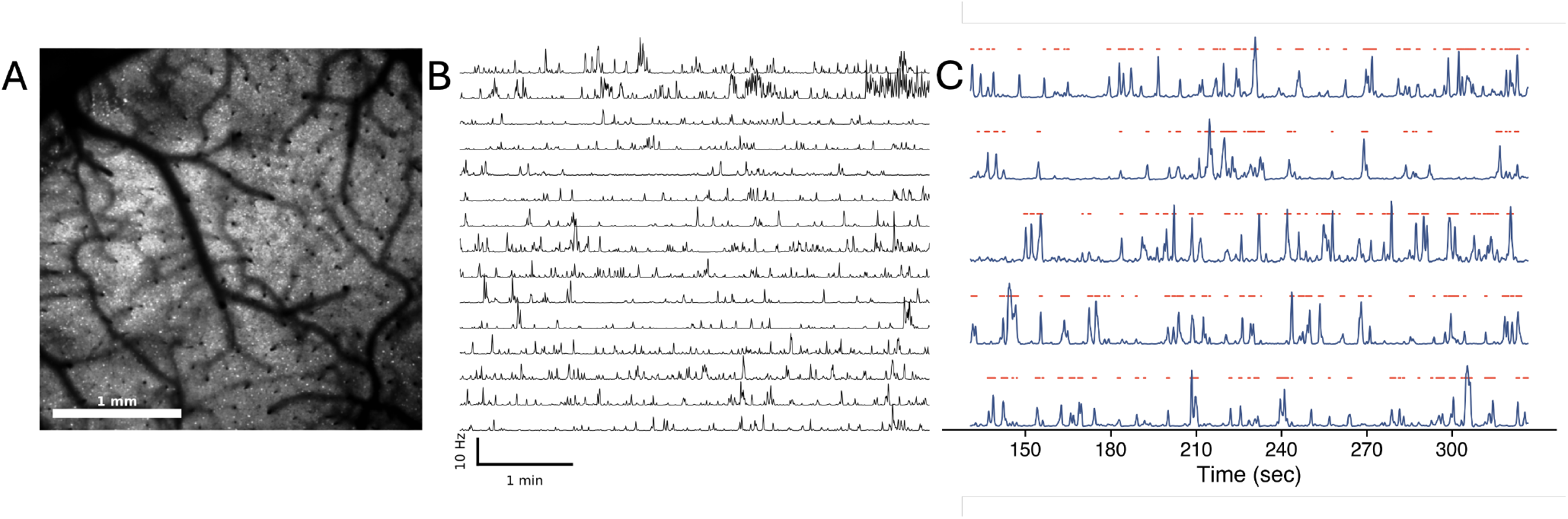
Data Preprocessing Pipeline. A) Recorded field of view from the Diesel2p mesoscope. Neurons are visible as the bright spots in the image. B) The deconvolved traces of spiking activity extracted from ROIs detected by the EXTRACT algorithm. C) The binarised states used for subsequent analysis. The blue traces represent the spiking activity of individual neurons, and the red dots indicate the on states detected by the Hidden Markov Model (HMM).

### 2.3 Information-Theoretic Measures

To measure the interdependencies between the neural activity traces recorded from the neurons in the visual cortex, we build upon mutual information (MI). MI quantifies the mutual dependence between two variables by computing the amount of information obtained about one random variable through observing another random variable. For two discrete random vectors **U** and **V**, the mutual information of **U** given **V**) is defined as:

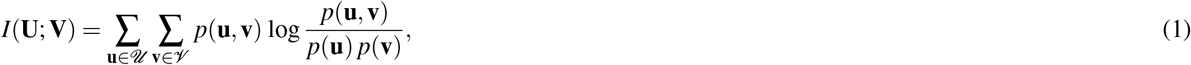

where **u** and **v** are (vector) values of the random variables **U** and **V**, respectively; *p*(**u, v**) is the joint probability distribution function; and *p*(**u**) and *p*(**v**) are the marginal probability distribution functions.

Here, we compute the mutual information between binarised and deconvolved calcium traces. The HMM-based binarisation helps to mitigate the effects of noise and variability in the calcium imaging data, and has been useful in the identification of neuronal assemblies^51^ and decoding behavior^52^.

#### 2.3.1 Partial Information Decomposition

Let us consider two neurons X and Y, with observable states *X*_*t*_ and *Y*_*t*_, respectively. To quantify the total information shared by this pair of neurons about their joint future activity after a time lag τ, we use the Time Delayed Mutual Information (TDMI), defined as *I*(*X*_*t*_,*Y*_*t*_; *X*_*t*+τ_,*Y*_*t*+τ_), where, according to the definition (1), we have **U** = [*X*_*t*_,*Y*_*t*_] given by the joint states of neurons X and Y at time *t* and **V** = [*X*_*t*+τ_,*Y*_*t*+τ_] by their joint states at a future time point *t* ++τ.

Using Partial Information Decomposition (PID)^33^, we can decompose the TDMI into four non-negative components (see Figure 2): unique information from neuron X (*UI*(*X*)), unique information from neuron Y (*UI*(*Y*)), redundant information (*RI*), and synergistic information (*SI*). The decomposition is given by:

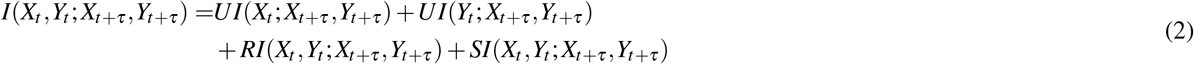

**Figure 2.**
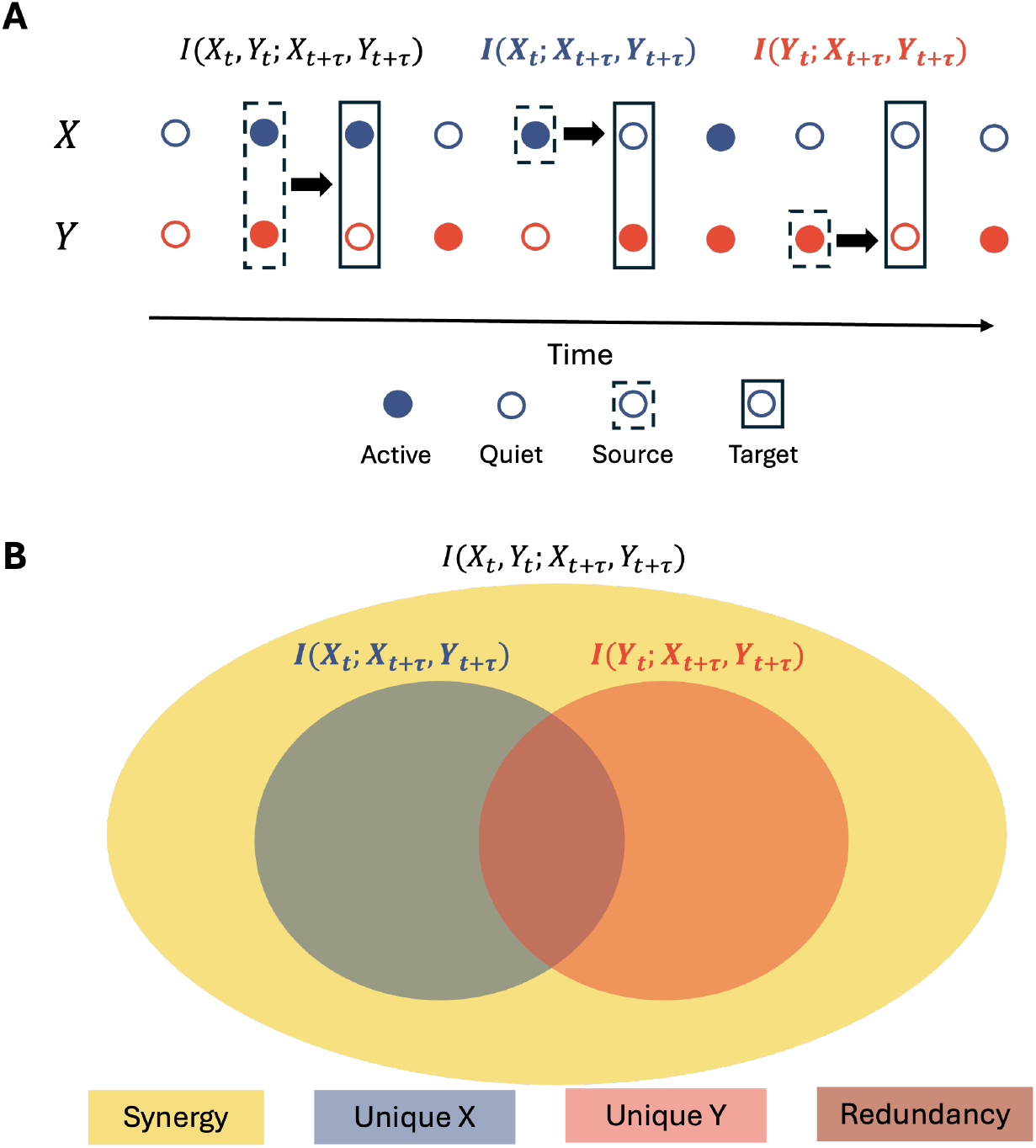
Partial Information Decomposition (PID) Framework. A) Schematic representation of two binary neurons X and Y, with their past states (*X*_*t*_, *Y*_*t*_) influencing their future states (*X*_*t*+τ_, *Y*_*t*+τ_) at a time lag τ. B) The TDMI between the past states of neurons X and Y (*X*_*t*_, *Y*_*t*_) and their joint future states (*X*_*t*+τ_, *Y*_*t*+τ_) can be decomposed into four non-negative components: unique information from neuron X (*UI*(*X*)), unique information from neuron Y (*UI*(*Y*)), redundant information (*RI*), and synergistic information (*SI*).

However, the PID framework does not provide a unique decomposition, as there are multiple ways to define the components. In this study, we employ the Minimum Mutual Information (MMI) approach to define the redundant information^53^ as:

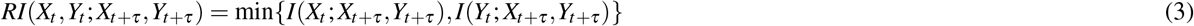

The MMI redundancy function provides an upper bound on the redundant information, but provides a non-negative and interpretable decomposition of the TDMI. The unique and synergistic information components can then be derived from the TDMI and the redundancy using the following equations:

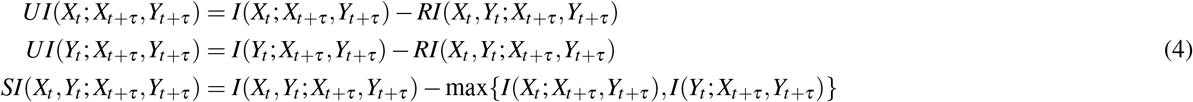

Here, *I*(*X*_*t*_; *X*_*t*+τ_,*Y*_*t*+τ_) and *I*(*Y*_*t*_; *X*_*t*+τ_,*Y*_*t*+τ_) are the mutual information between the past state of neuron X (or Y) and the joint future state of both neurons, [*X*_*t*+τ_,*Y*_*t*+τ_]. Note that, by the definition (3), one of the unique information components will be zero, depending on which neuron has the lower mutual information with the joint future state. In our analysis, we restrict ourselves to τ =1 as the time delay.

#### 2.3.2 Null Models for Information-Theoretic Measures (NuMIT)

Our study involves the comparison of information-theoretic measures both within a dataset (across different recording paths) and across different datasets. Therefore, it is crucial to normalise these measures to account for potential biases arising from finite data size, firing rates, redundancy functions, and other confounding factors. To address this, we employ a null model-based normalisation approach called NuMIT (Null Models for Information-Theoretic Measures)^54^.

NuMIT provides a systematic way to normalise the decomposed PID components by comparing them to null models that exhibit the same TDMI as the experimental data. The null models are generated by simulating a pair of neurons using randomly sampled binary processes constrained by adding independent noise to match the experimental TDMI. This ensures that the null models capture the same level of overall information transfer as the empirical data, while allowing us to assess the significance of the individual PID components. The normalisation is performed by calculating the Z-scores of the empirical PID components relative to the null distributions. We can thus identify which components are significantly different from the expected value of the null models, constrained so that the overall time delayed information is preserved. The NuMIT framework has been shown to mitigate biases in information-theoretic measures and to accurately assess the underlying information dynamics in neural systems^54^.

#### 2.4 Correlation Length

To quantify the correlation length in the neural activity, we compute the pairwise Pearson correlation coefficient between the binarised spiking activity of all pairs of neurons in each dataset. The correlation coefficients are binned according to the inter-neuron distance, and the average correlation coefficient is then computed for each of the logarithmically spaced distance bins. The correlation length *λ* is then estimated by fitting an exponential decay of the average correlation coefficient with distance:

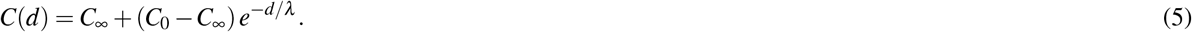

Here, *C*(*d*) is the average correlation coefficient at distance *d, C*_0_ is the initial correlation coefficient at the minimum distance, *C*_∞_ is the asymptotic correlation coefficient at large distances, which accounts for the baseline correlation in mesoscale calcium imaging data at large distances. The correlation length *λ* provides a characteristic length scale for the decay of correlations in space. The parameters, *C*_0_, *C*_∞_ and *λ* in equation 5 were estimated by a non-linear least squares fitting algorithm using the *SciPy* library in Python^55^. To visualise the decay of correlations with distance, we also defined the normalised average correlation coefficient:

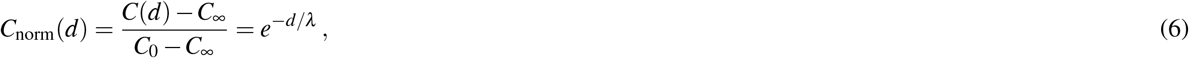

which allows for consistent comparison of the decay of correlations across different datasets and conditions.

The Z-scored Synergy and Redundancy values decay slowly with distance, with fitted *λ* beyond the field of view. To obtain more robust estimates of the spatial extent of information decay in the neural activity, we estimate the *effective information length λ*_eff_ as the normalised area under the curve of the fitted exponential decay function:

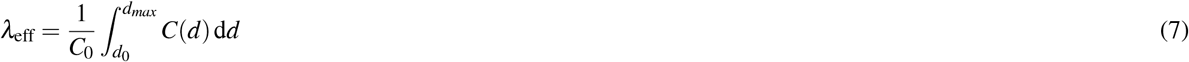

where *d*_0_ = 100*μm* is the minimum distance and *d*_*max*_ = 1500*μm* is the maximum distance in the field of view such that sufficient pairs of neurons are available.

#### 2.4.1 Partial Network Decomposition

Within each dataset, we compute the pairwise synergistic and redundant interactions between neurons based on the normalised PID components. We formalise these interactions as networks, where the nodes are neurons and the edge weights correspond to the normalised synergistic (or redundant) information between the neurons. We then obtain the associated k-nearest neighbour (kNN) graph, whereby we retain only the top *k* strongest connections for each neuron. This results in a sparsified network representation that highlights the most relevant interactions while reducing noise and spurious connections.

To carry out our analysis for the partial network decomposition, we consider the two unweighted networks (synergistic and redundant) and a *combined network* that contains the union of the edge sets from the synergistic and redundant layers (see Figure 3). This combined representation allows us to analyse the interplay between synergistic and redundant interactions and their contributions to the overall information propagation in the neural system. Using partial network decomposition^56^, we identify complementary, shared, and unique shortest paths between pairs of neurons across the synergistic and redundant layers (see Figure 3). This analysis provides insights into how the different types of information interactions contribute to the overall connectivity and information flow in the network across different spatial scales.

**Figure 3.**
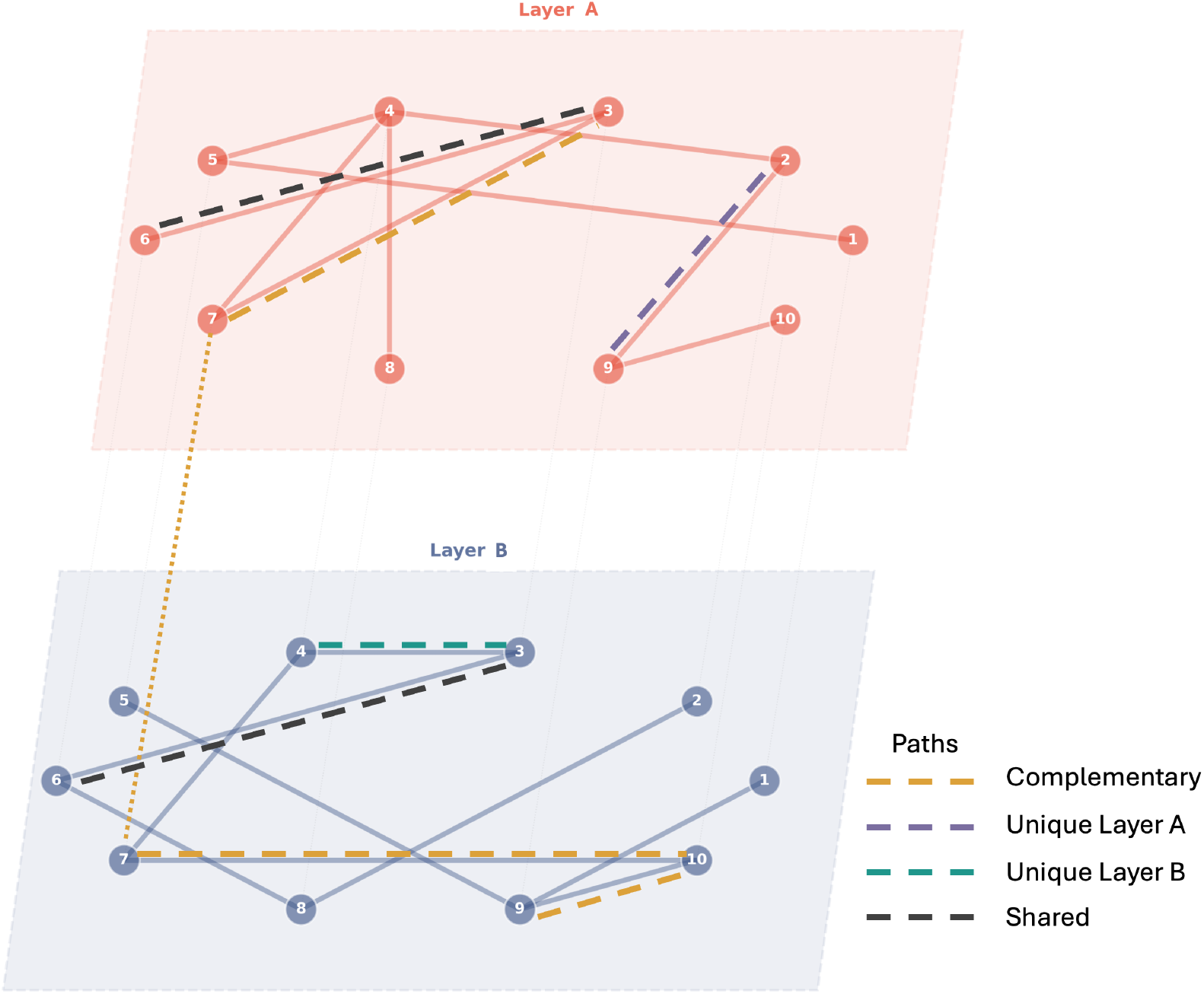
Partial Network Decomposition. Schematic representation of a combined network formed by Layers A and B. The different types of shortest paths between the nodes are illustrated using dashed lines: Complementary Paths (orange), Shared Paths (black), and Unique Paths (purple and teal). For instance, the shortest path between nodes 3 and 9 is complementary (orange), as it leverages connections from both layers to achieve a shorter path than either layer alone. The edge between nodes 3 and 6 is present in both layers, representing a shared path (black). The shortest path between nodes 2 and 9 is unique to Layer A (purple), while the shortest path between nodes 3 and 4 is unique to Layer B (teal).

Briefly, the framework considers shortest paths between pairs of nodes as a measure of efficiency. Suppose we have a combined network with two layers, A and B, representing synergistic and redundant interactions, respectively. For each pair of nodes (*i, j*), we identify the shortest paths in each layer separately, denoted as *d*_*A*_(*i, j*) and *d*_*B*_(*i, j*). We then consider the shortest path on the combined network, which includes the union of the edges from both layers, and we denote it as *d*_*A∪B*_(*i, j*). The partial network decomposition then classifies the shortest paths into three categories:

- **Complementary Paths**: *d*_*A∪B*_(*i, j*) *<* min(*d*_*A*_(*i, j*), *d*_*B*_(*i, j*)). This indicates that the interaction between the two layers provides a more efficient route for information transfer than either layer alone.
- **Shared Paths**: *d*_*A∪B*_(*i, j*) = max(*d*_*A*_(*i, j*), *d*_*B*_(*i, j*)). This indicates that the interaction between the two layers does not provide any additional efficiency for information transfer beyond what is already available in either layer.
- **Unique Paths**: *d*_*A∪B*_(*i, j*) = min(*d*_*A*_(*i, j*), *d*_*B*_(*i, j*)) and *d*_*A∪B*_(*i, j*) *<* max(*d*_*A*_(*i, j*), *d*_*B*_(*i, j*)). This indicates that one layer provides a more efficient route for information transfer than the other layer, and the interaction between the two layers does not provide any additional efficiency.

By analysing the distribution of complementary, shared, and unique paths, we can characterise how the contribution of synergistic and redundant interactions varies over paths of different lengths in the network.

## 3 Results

### 3.1 Long-Range Correlations in the Visual Cortex

We first investigate the presence of long-range correlations in the neural activity recorded from the visual cortex of awake mice. In Figure 4A (left), we plot the average Pearson correlation coefficients between the binarised spiking activity of two neurons as a function of the distance between the neurons (binned), for both spontaneous and visually stimulated conditions. The results show that the correlations decay with distance, but remain significantly above zero even at distances of several millimetres, indicating the presence of long-range correlations in the neural activity. The fitted correlation functions (Eq. (5)) are also shown in Figure 4A (left). From each fit, we obtain values for the initial correlation *C*_0_, correlation length *λ* and asymptotic correlation *C*_∞_. The normalised correlations (Eq. (6)) shown in Figure 4A (right) highlight the increased correlation length observed during visual stimulation. Figure 4B shows that the initial correlation *C*_0_ is not significantly different (Mean difference = 0.002, *p* = 0.0871 and Hedge’s *g* = 0.91) among the datasets in the two conditions, yet the estimated correlation lengths are significantly larger (Mean difference = 603.21*μm, p* = 0.0055 and Hedge’s *g* = 2.25) during visual stimulation compared to spontaneous activity, suggesting that visual input enhances long-range correlations in the visual cortex. An increased correlation length is a signature of a system operating closer to criticality^32^. It must be noted that although an increase in correlations is expected due to the visual stimulus for short distances (*≈* 200 −300*μm*)^57^, we observe increased correlations up to ≈ 900*μm*. This indicates a distinct state of the cortical network that supports longer-range correlations rather than short-range stimulus-related co-activations.

**Figure 4.**
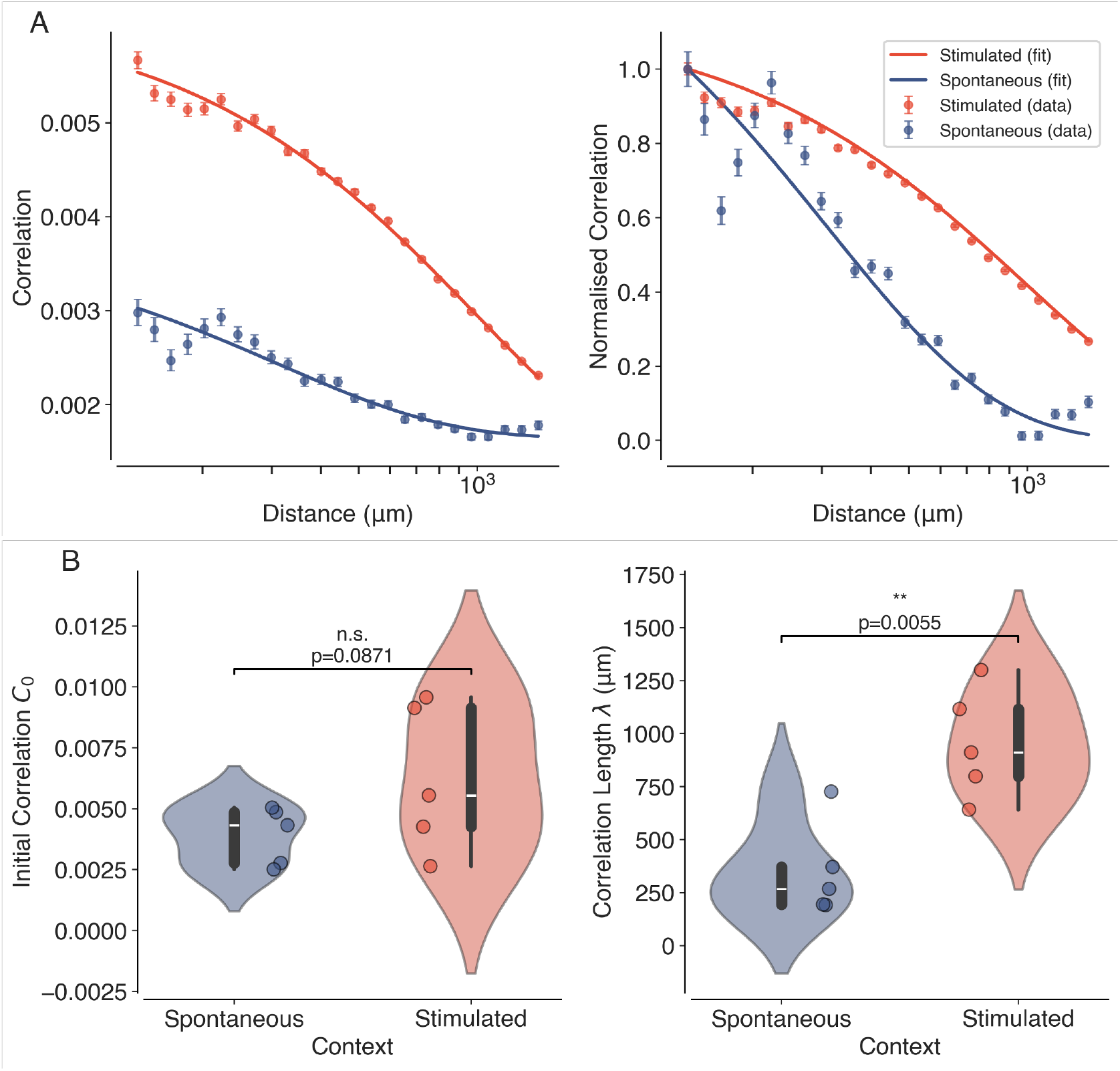
Long-Range Correlations in the Visual Cortex. (A) Average Pearson correlation coefficients (left) as a function of distance between pairs of neurons, for both spontaneous (blue) and visually stimulated (orange) conditions. The normalised correlation coefficients are shown on the right. The solid lines represent the fitted exponential decay functions. The slower decay of correlations in the stimulated condition is clearly visible in the normalised plots. (B) Violin plots showing the estimated correlation lengths *λ* for spontaneous and visually stimulated conditions across all datasets. Individual data points (*N* = 10, 5 Spontaneous and 5 Stimulated) are overlaid as dots. Asterisks indicate statistical significance (^∗∗^, *p <* 0.01), estimated using permutation tests.

### 3.2 Information Decomposition Reveals Distinct Spatial Profiles of Synergy and Redundancy

Next, we apply PID and NuMIT to decompose the time-delayed mutual information (TDMI) between pairs of neurons into Z-scored redundant and synergistic components. The TDMI was estimated at a time delay of τ = 1 time step. The average Z-scored synergy and redundancy as a function of distance between pairs of neurons are shown in Figure 5. Both redundancy (Fig. 5A, left) and synergy (Fig. 5B, left) exhibit a decay with distance, but average initial redundancy (Z-score = 1.24*±* 0.06) is generally higher than synergy (Z-score = 0.23*±* 0.021), and the decay is slower for redundancy (*λ*_eff_ = 1278 *±*11*μm*) compared to synergy (*λ*_eff_ = 1230 *±μm*). This suggests that redundant information is more prevalent and more spatially extended in the visual cortex. These findings are consistent with previous studies that have reported that sensory regions of the brain exhibit high levels of redundancy, which may serve to enhance the robustness and reliability of sensory processing^58^. The normalised Z-score plots on the right highlight the distinct spatial profiles of synergy and redundancy in the two contexts (see Figure 5 A, B, right). While the spatial decay of redundancy is similar in both spontaneous and visually stimulated conditions, synergy exhibits a slower decay during visual stimulation compared to spontaneous activity. This indicates the enhanced role of synergistic interactions in supporting long-range correlations during visual processing.

**Figure 5.**
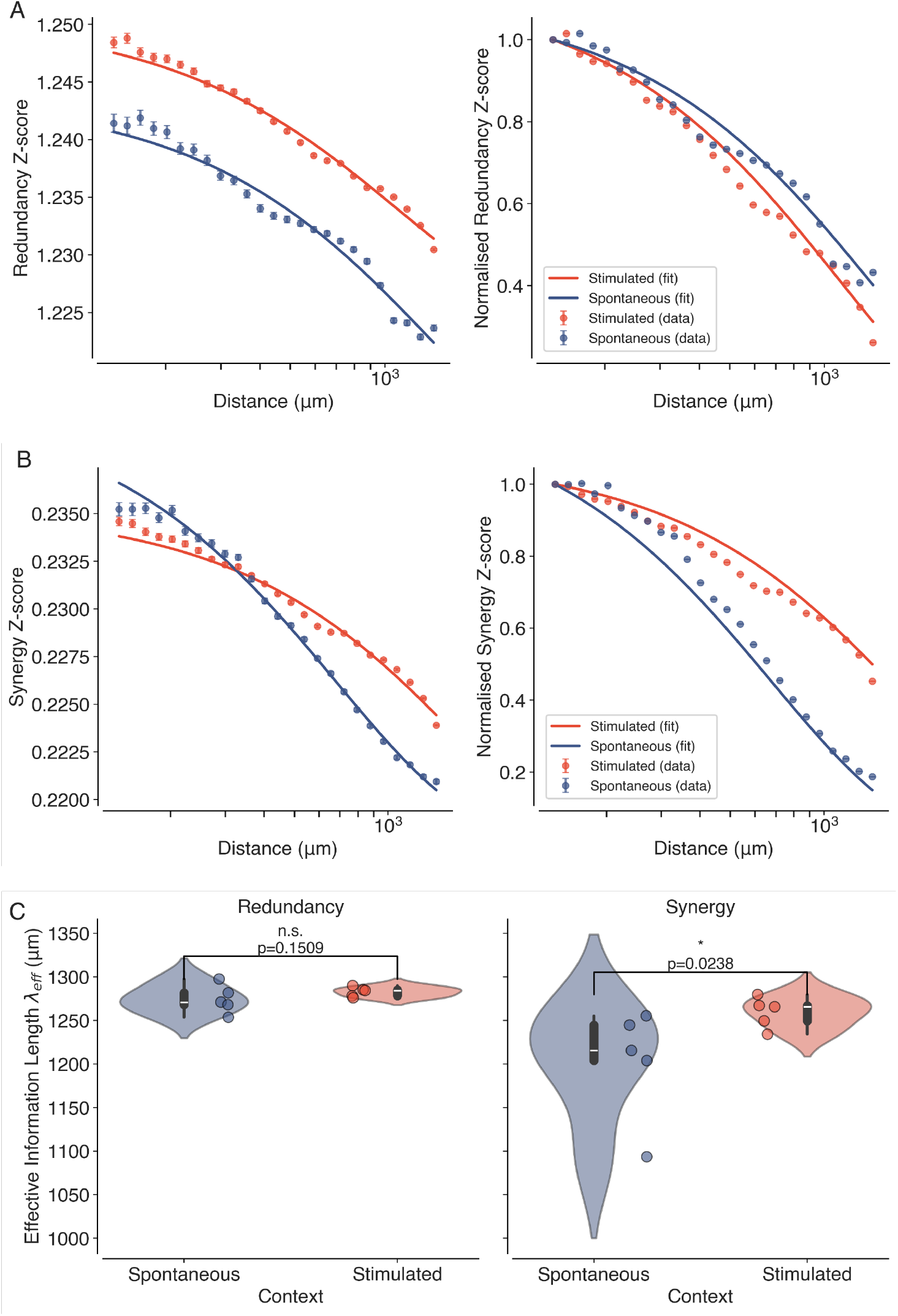
Information Decomposition Reveals Distinct Spatial Profiles of Synergy and Redundancy. (A) Average Z-scored synergy and redundancy values as a function of distance between pairs of neurons, for both spontaneous (blue) and visually stimulated (orange) conditions. The solid lines represent the fitted exponential decay functions (Eq. 5). (B) Violin plots showing the estimated effective correlation lengths *λ*_eff_ for synergy and redundancy across all datasets (*N* = 10, 5 Spontaneous and 5 Stimulated) in spontaneous (blue) and visually stimulated conditions (orange). Individual data points are overlaid as dots. Asterisk indicates statistical significance (^∗^, *p <* 0.05), estimated using permutation tests.

The fits also allow us to obtain *effective information lengths* for synergy and redundancy across all datasets, as shown in Figure 5C. We observe that while the redundancy information lengths do not significantly change (Mean difference = 8.33*μm, p* = 0.150 and Hedge’s *g* = 0.61) between spontaneous and stimulated datasets, the synergy information lengths are significantly larger (Mean difference = 56.67*μm, p* = 0.023 and Hedge’s *g* = 1.08) during visual stimulation compared to spontaneous activity. This suggests that visual stimulation enhances the spatial extent of synergistic interactions in the visual cortex, which may facilitate more efficient information processing and integration across different regions, whereas stimulation has little effect on the spatial extent of redundant interactions. Overall, these results highlight the unique role of synergistic interactions in mediating the increase of the spatial extent of long-range correlations, a signature of criticality.

### 3.3 Partial Network Decomposition Reveals Complementary Roles of Synergy and Redundancy Networks

To further investigate the interplay between synergistic and redundant interactions in the visual cortex, we apply partial network decomposition to combined networks, where neurons are nodes and the unweighted edges of the sparsified redundancy and synergy layers are constructed from the corresponding normalised components.

We then compute the complementary, shared, and unique paths that contribute to the propagation of information in the combined network representing the neurons of the visual cortex and their different interactions.

The proportion of complementary, shared, and unique paths as a function of path length is shown for both the spontaneous and stimulated conditions in Figure 6. We observe that the proportion of complementary paths increases with path length, indicating that synergistic and redundant interactions work together to facilitate efficient information processing across longer distances. The proportion of unique paths decreases with path length, and there are very few shared paths across all path lengths. These findings suggest that synergistic and redundant interactions complement each other over longer paths while maintaining a unique presence over shorter paths.

**Figure 6.**
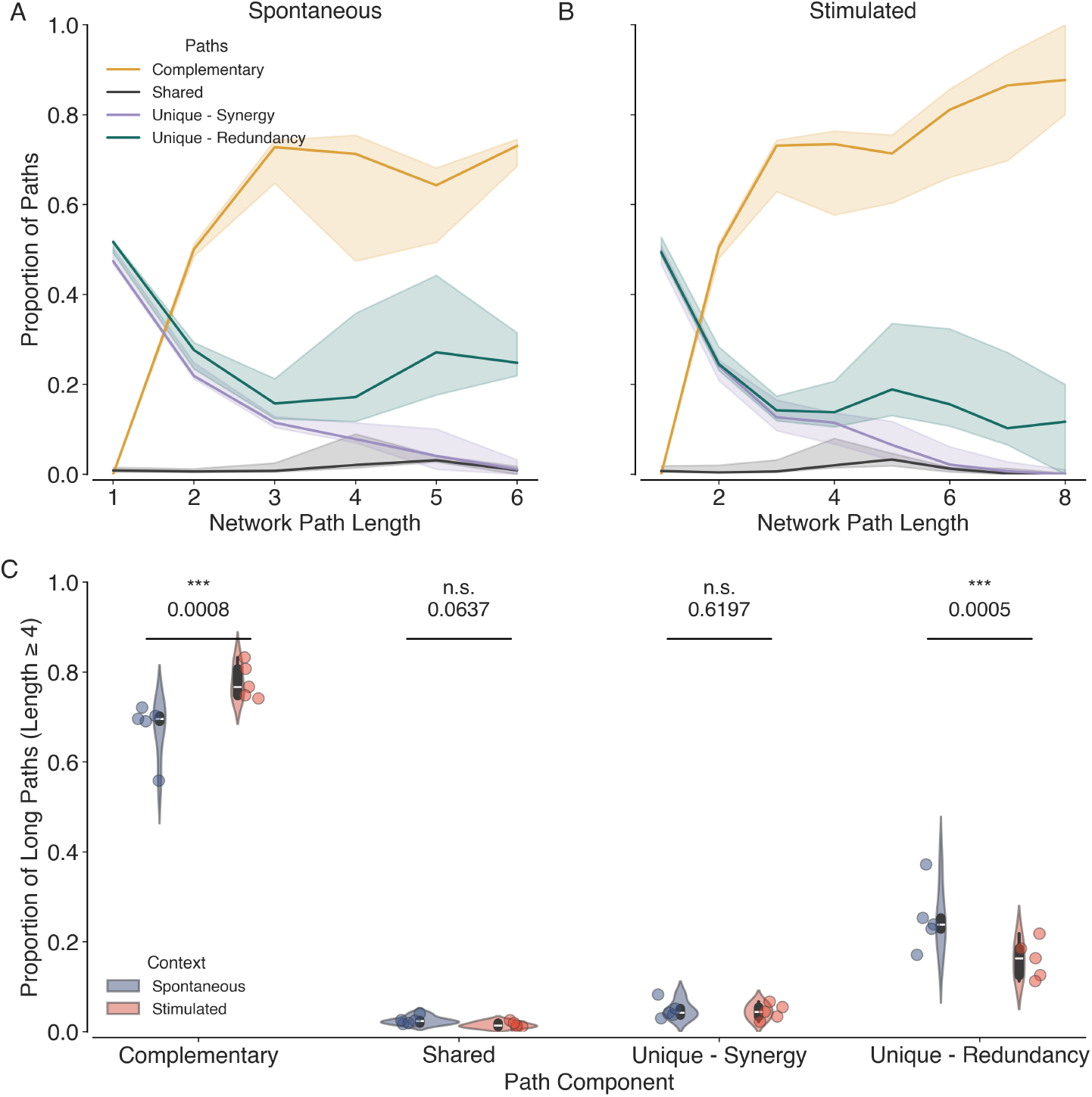
Partial Network Decomposition Reveals Complementary Roles of Synergy and Redundancy networks. Proportion of complementary (orange), shared (black), and unique paths (purple and teal) as a function of network path length for both spontaneous (A) and stimulated (B) conditions. The solid lines represent the median across all datasets, and the shaded areas represent 95% confidence intervals. (C) Violin plots showing the proportion of long complementary, shared and unique paths among the synergy and redundancy layers across all datasets (*N* = 10, 5 Spontaneous and 5 Stimulated) in spontaneous (blue) and stimulated conditions (orange). Individual data points are overlaid as dots. Asterisk indicates statistical significance (^∗∗∗^, *p <* 0.001) estimated via permutation tests.

During stimulation, we observe a significant increase in the proportion of long (path length ≥4) complementary paths (Mean difference = 0.105, *p* = 0.0008) and a decrease in the proportion of long unique paths in the redundancy layer (Mean difference = −0.09, *p* = 0.0005), compared to spontaneous activity. This suggests that enhanced cooperative interactions between synergy and redundancy networks are observed at the expense of unique paths in the redundancy network during visual stimulation. These results highlight the dynamic nature of information interactions in the brain and their modulation by sensory input.

## 4 Discussion

In this study, we investigated the presence of long-range correlations in the neural activity recorded from the visual cortex of awake mice and explored the role of synergistic and redundant interactions in mediating these correlations. Using two-photon calcium imaging data from the mesoscope, we found that the visual cortex exhibits long-range correlations that are enhanced during visual stimulation. By applying the PID framework, we decomposed the time-delayed mutual information into synergistic and redundant components, and found that synergy plays a crucial role in mediating long-range correlations. Furthermore, the partial network decomposition revealed that both synergistic and redundant interactions cooperatively enable information processing over long distances in the visual cortex, especially under stimulation, when complementary paths become more important at the expense of unique redundant paths.

By identifying the presence of long-range correlations in the visual cortex, we provide support for the brain criticality hypothesis, which posits that the brain operates near a critical point to optimally process information and respond to a wide range of stimuli^10–12^. The dilated correlation lengths observed during visual stimulation suggest that sensory input can modulate the critical state of the brain, potentially allowing for more efficient information processing and integration across different regions. However, it must be noted that increased correlation lengths can arise due to increased arousal^59^ or attention^60^, and not necessarily due to the visual stimulation itself. In future work, we are exploring the differential roles of arousal and stimulus in explaining the move towards criticality in the visual cortex.

We observe that redundant interactions are stronger and decay more slowly than synergistic interactions in the primary visual cortex. This finding is in line with the view of redundancy as a mechanism for enhancing the robustness of information processing^58^. On the other hand, synergistic interactions exhibit a more pronounced increase at large distances during visual stimulation, suggesting their unique role in coordinating spatially distributed information processing. We note that this selective enhancement of synergy departs from the increase in both synergy and redundancy near criticality in traditional models such as the Ising model^61^. This suggests that the brain may employ more complex mechanisms to regulate information interactions, beyond what is captured by simple models of criticality. Future studies could explore which biological mechanisms can explain the selective enhancement of synergy when approaching criticality in computational models of brain criticality.

Our work highlights the importance of considering different types of information interactions in understanding neural systems. Although this study focuses on time-delayed pairwise interactions, future studies could extend this framework to higher-order (beyond pairwise) interaction measures. Furthermore, although our focus has been on the primary visual cortex, it would be interesting to explore how these findings generalise to other brain regions and cognitive processes. The tools and frameworks presented here provide an approach to study the link between long-range correlations and information propagation mechanisms in neural systems, and could be applied to other datasets and experimental paradigms.

## Author Contributions

All authors contributed to the review and editing of the manuscript. HR and HJJ conceptualised the study. JC performed the experiments and data acquisition. CS and MGK preprocessed the calcium imaging data. HR analysed the data and wrote the initial draft of the manuscript. MS conducted the network analysis. MB, SS, SLS and HJJ supervised the project. All authors contributed to the final draft of the article and approved the submitted version.

## Acknowledgments

We thank the members of the Statistical Physics of Cognition project, Prof. Lucilla de Arcangelis, Dr. Pedro Mediano, Dr. Fernando Rosas and Alberto Liardi for helpful discussions and feedback on the manuscript. We also thank the Imperial College High Performance Computing Service for computational resources.

